# Small cationic cysteine-rich defensin-derived antifungal peptide controls white mold in soybean

**DOI:** 10.1101/2023.05.16.540985

**Authors:** Arnaud Thierry Djami-Tchatchou, Meenakshi Tetorya, Jennette M. Codjoe, Dilip M. Shah

## Abstract

White mold disease caused by a necrotrophic ascomycete pathogen *Sclerotinia sclerotiorum* results in serious economic losses of soybean yield in the USA. Lack of effective genetic resistance to this disease in soybean germplasm and increasing pathogen resistance to fungicides makes white mold difficult to manage. Small cysteine-rich antifungal peptides with multi-faceted modes of action hold potential for development as sustainable spray-on bio-fungicides. We have previously reported that GMA4CG_V6 peptide, a 17-amino acid variant of the MtDef4 defensin-derived peptide GMA4CG containing the active γ-core motif, exhibits potent antifungal activity against the gray mold fungal pathogen *Botrytis cinerea in vitro* and *in planta*. This peptide exhibited antifungal activity against an aggressive field isolate of *S. sclerotiorum 555 in vitro.* It markedly reduced white mold disease symptoms when applied to detached soybean leaves, pods, and stems. Spray-application on soybean plants provided robust control of the white mold disease. GMA4CG_V6 at sub-lethal concentrations reduced sclerotia production. It was also non-phytotoxic to soybean plants. Our results demonstrate that GMA4CG_V6 peptide has high potential for development as a bio-fungicide for white mold control in soybean.

## 1. Introduction

Soybean (*Glycine max* L. Merrill) is the second largest agricultural crop grown in the USA. In 2021, its production reached a value of $57.5 billion [1]. However, soybean production is severely impacted by white mold disease which ranks in the top 10 most destructive diseases of soybeans [2]. White mold is caused by *Sclerotinia sclerotiorum* (Lib.) de Bary, a necrotrophic fungal pathogen capable of infecting over 500 plant species worldwide. *S. sclerotiorum* can infect young seedlings, mature plants and pods at the pre- and post-harvest stage [3, 4]. It can persist for years in the field due to the long-term survival of hard resting structures called sclerotia, formed by the aggregation of fungal hyphae [5]. The sclerotia can survive in soil for many years and infect host plants by producing ascospores or mycelia [3, 6]. In soybean only partially resistant varieties are available to growers [7, 8].

White mold is currently managed through cultural practices, biological control and chemical fungicides [3, 9]. Its control relies mainly on application of fungicides that have single-site mode of action (MoA). However, fungicide resistance in pathogen populations is rising, rendering fungicide applications ineffective [10, 11]. There is an urgent need to develop safe and sustainable multi-target biofungicides to counter fungicide resistance.

Plants express a large number of diverse, small cysteine-rich antimicrobial peptides (AMPs) which exhibit potent antifungal activity and act as key components of their innate immune system. AMPs provide a first line of defense against invading pathogens [12-14]. Defensins with tetradisulfide arrays comprise a major class of AMPs in the plant kingdom. They share structural similarity but are diverse in their primary sequences. These cationic peptides exhibit broad-spectrum antifungal activity in a low micromolar or submicromolar range and have multi-faceted MoA [15, 16]. Major determinants of the antifungal activity of plant defensins reside in their γ-core motif (GXCX_3-9_C) composed of β2-β3 strands and an interposed loop. Artificial defensin-derived peptides containing γ-core motifs possess potent antifungal activity against fungal pathogens and are attractive candidates for commercial development as spray-on bio-fungicides to replace conventional chemical fungicides [17, 18].

To date only one defensin, RsAFP1, has been reported to have antifungal activity *in vitro* against *S. sclerotiorum* at 20 µg/ml (3.5 µM) [19]. In contrast, multiple AMPs with antifungal activity at low micromolar concentrations *in vitro* against necrotrophic gray mold pathogen *Botrytis cinerea* have been reported [20-22]. Both *S. sclerotiorum* and *B. cinerea* belong to the family *Sclerotiniaceae*, share genome sizes of 38-39 Mb, and have similar genes governing necrotrophy [23].

GMA4CG_V6 is a synthetic peptide with the sequence GGRCKGFRRRWFWTRIC. It is a variant of the GMA4CG peptide comprising the last 17-amino acids of the plant defensin MtDef4 [21]. It contains a functionally active γ-core motif, one C3-C17 disulfide bond, and five amino acid substitutions. It was recently characterized for its *in vitro* and *in planta* spray-on fungicidal activity against *B. cinerea* [21]. In the present study, we found that GMA4CG_V6 also has antifungal activity against *S. sclerotiorum 555*. It exhibits fungicidal activity against this pathogen *in vitro,* on detached soybean leaves, pods and stems and also when applied topically on tobacco and soybean leaves *in planta*. At sub-lethal concentrations, GMA4CG_V6 inhibits sclerotia production. Thus, GMA4CG_V6 has potential for development as a multi-target bio-fungicide for control of white mold.

## 2. Materials and Methods

### 2.1. Plant and Fungal Materials

Williams 82 soybean plants (*Glycine max* L. Merrill) were used in this study. Plants were grown in soil in a greenhouse with a short-day photoperiod (8h light/16h dark) at 21°C and 75% relative humidity, with a light intensity of ∼130 μEinsteins sec^-1^ m^-2^. They were inoculated at approximately 3-4 weeks of age. A highly aggressive strain of *S. sclerotiorum 555* on soybean was used for this study [10]. This strain and strains *1902* and *1922* were kindly provided by Dr. Sydney Everhart of the University of Nebraska, USA. Fungal cultures were grown on potato dextrose agar (PDA) medium (Difco Laboratories Inc., Detroit, MI, USA) at room temperature.

### 2.2 In vitro Antifungal Activity of GMA4CG_V6 against S. sclerotiorum

The *in vitro* antifungal activity of GMA4CG_V6 against *S. sclerotiorum 555*, *1902* and *1922* was determined using a 24-well plate assay. Mycelial plugs (1 mm in diameter) cut from the growing edge of the two day-old actively growing colony were transferred from PDA media to a series of wells containing 250 μL of 1x synthetic low-salt fungal medium (SFM) [27] and GMA4CG_V6 at final concentrations of 3, 6, 12, 24 and 48 µM or water. The plates were incubated at room temperature for 2 days prior to assessment. The diameter of mycelial growth at 2 dpi was measured using ImageJ software. The inhibition of fungal growth was calculated as a percentage using the formula: IG (%) = (*D*c – *D*t)*/D*c × 100; where IG is inhibition of growth, *D*c is the mycelium diameter (mm) of the control and *D*t is the mycelium diameter (mm) of the treated dishes. These experiments were performed three times.

### 2.3. Semi-in planta Antifungal Activity of GMA4CG_V6 against S. sclerotiorum 555

For the semi-*in planta* antifungal assays, a 1 mm mycelial plug was taken from the leading edge of a 2-day old fungal colony growing on PDA media and placed on detached leaves of three to four-week-old Williams 82 plants or pods and stems of approximately six-week-old plants. Then, 40 μL of GMA4CG_V6 at 6, 12, 24, 48 and 96 µM or H_2_O (control) was applied immediately on the plug. Inoculated leaves, pods, and stems were allowed to incubate under high humidity in the dark for 2-7 days prior to assessment. Lesions were photographed and the area of fungal lesion areas at 2dpi were measured using ImageJ software. The severity of disease lesions on each leaf was assessed using the CropReporter system as described in [22]. The quantification of plant health and stress was carried out using the calculated F_V_/F_M_ (maximum quantum yield of photosystem II) images of the CropReporter system (PhenoVation, Wageningen, Netherlands), showing the efficiency of photosynthesis in false colors. These experiments were performed three times.

### 2.4 In planta Antifungal Activity of GMA4CG_V6 against S. sclerotiorum 555

The antifungal activity of GMA4CG_V6 against *S. sclerotiorum 555* was tested *in planta* using 3-week-old *Nicotiana benthamiana* plants or 3 to 4-week-old Williams 82 soybean plants. A suspension of ground mycelium (1 mL) in potato dextrose broth (PDB) at optical density (OD) of 0.5 was used to spray leaves followed by the spray of 2 mL of different concentrations of GMA4CG_V6 or water. Then, the *in planta* antifungal activity of GMA4CG_V6 against *S. sclerotiorum 555* was also tested against 3 to 4-week-old Williams 82 soybean plants. The pots were incubated under high humidity for 2 days prior to assessment. Fungal lesion area at 2 dpi was measured using ImageJ software and high-resolution fluorescence images were taken using CropReporter (PhenoVation, Wageningen, Netherlands) and analyzed as described above.

### 2.5 . Effect of GMA4CG_V6 on Sclerotia Production in S. sclerotiorum 555

We explored the effect of GMA4CG_V6, GMA4CG_V6_lin without a disulfide bond and the GMA4CG_V6_Ala3 variant carrying the substitution of RRRW with AAAA (Tetorya et al, 2023) on sclerotial development in *S. sclerotiorum*. A previous study showed that the formation of sclerotia on PDA plates starts at the 4^th^ day of the mycelial growth and ends at the 9^th^ day when they have a black coloration and are easy to detach from the culture medium [28]. To assess the effect of GMA4CG_V6 on sclerotia production, mycelial plugs cut from the margin of a 2-day-old colony were transferred onto PDA plates and allowed to grow for one day. Since we found that GMA4CG_V6 was inactive when amended in PDA medium, 500 µl of 12 µM GMA4CG_V6 was sprayed on the mycelia at 1^st^, 3^rd^ and 5^th^ day. PDA plates sprayed only with water were used as controls. After treatment, the PDA plates were incubated at 25°C for nine days in the dark and then the production of sclerotia was evaluated. The experiment was performed in quadruplicate for each treatment with similar results.

### 2.6 Effect of GMA4CG_V6 on the Expression of S. sclerotiorum Genes Related to Sclerotia Production

It has been previously shown that expression of genes involved in sclerotial development-calcineurin *(cna1) (SS1G_01788.1)*, transcriptional regulatory protein *(Pac1) (SS1G_07335.1),* Sch9-like protein kinase (*Pka2) (SS1G_01124.1)* and MAP kinase (*Smk1) (SS1G_11866.1)*-is significantly reduced in *S. sclerotiorum* when the inhibitory compound cinnamic acid is applied at its EC_50_ value [24, 29]. To monitor the effect of GMA4CG_V6, GMA4CG_V6_lin and GMA4CG_V6_Ala3 on the expression of these genes, mycelial plugs (5 mm) were used to inoculate PDB media containing water (untreated control) or cinnamic acid (25 µg/ml) as our positive control or each peptide (12 µM) in a total volume of 1 ml. Samples were incubated at 26 °C with shaking at 160 rpm for two days. After two days, mycelia were collected from PDB and flash-frozen in liquid nitrogen; then total RNA was extracted using the Nucleospin RNA Isolation Kit (Katara Bio, San Jose, CA, USA). The RNA was reverse transcribed to first strand cDNA using the Revertaid™ cDNA synthesis kit (Thermo Scientific, Waltham, MA, USA). qPCR was then used to monitor the expression of these genes using ig™ SYBR® Green qPCR 2x master mix (Intact Genomics, St Louis, MO, USA) on a CFX Connect real-time PCR detection system (Bio-Rad, Hercules, CA, USA) following the manufacturer’s instructions. In each experiment, we used three biological replicates with two technical replicates for each. The relative expression was determined using the method of Pfaffi [30] and the qPCR data were normalized using actin (XM_001589919.1) as reference gene. The actin gene primers used are from the study of [24] and those for β-tubulin (MN296024.1) are from [31] (Table S1).

### 2.7. SYTOX Green^TM^ (SG) Plasma Membrane Permeabilization Assay

The effect of GMA4CG_V6 on the membrane integrity of *S. sclerotiorum* 555 was determined using a modified SG uptake quantification assay [32, 33]. Membrane permeabilization of fungal cells was analyzed using confocal microscopy by visualizing the influx of the fluorescent dye SG (Thermo-Fisher Scientific, Waltham, MA, USA). Fresh germlings mixed 12 μM GMA4CG_V6 and 1 μM SG were deposited onto glass-bottom petri dishes and were imaged by confocal microscopy at an excitation wavelength of 488 nm and an emission wavelength ranging from 520 to 600 nm at specific time intervals. Plates with SG but without GMA4CG_V6 were used as negative controls. A Leica SP8-X confocal microscope was used for all confocal imaging, and Leica LASX (version 3.1.5.) software was used to process these images.

### 2.8 Uptake of GMA4CG_V6 by S. sclerotiorum 555 Cells

Time-lapse confocal laser scanning microscopy was performed to monitor uptake of the Tetramethyl rhodamine (TMR)-labeled peptide as described previously [32]. Since labeled GMA4CG_V6 lost approximately 50% of its antifungal activity, it was used at a final sub-lethal concentration of 24 µM. The excitation and emission wavelength for TMR were 562 and 580 to 680 nm. The confocal microscope images was captured 2-5 min after GMA4CG_V6 challenge.

### 2.9. Assessment of the Effects of GMA4CG_V6 Treatment on Growth of Soybean Plants

To test the effects of GMA4CG_V6 on soybean plants, 48 µM of GMA4CG_V6 or water was sprayed on two-week-old soybean plants and the plants were grown in the growth chamber. For each treatment, five plants were used. Six weeks post-treatment, above ground portions of the plants were harvested, root systems were collected and washed of soil, and samples were oven dried at 60°C for 3 days and then weighed.

### 2.10. Statistical Analysis

Datasets were statistically compared with the statistical analysis software GraphPad Prism 8.0 (GraphPad software, San Diego, CA, U.S.A.), using one-way analysis of variance, followed by the Tukey’s post hoc test or Student’s t-test. The confidence level of all analyses was set at 95%, and values with *P* <0.05 were considered significant.

## 3. Results

### 3.1. In vitro Antifungal Activity of GMA4CG_V6 against S. sclerotiorum 555

We first tested the *in vitro* antifungal activity of GMA4CG_V6 against an aggressive isolate of *S. sclerotiorum* strain *555* [10]. We inoculated synthetic fungal medium (SFM) containing various concentrations of GMA4CG_V6 with 1 mm mycelial plugs of this pathogen. *S. sclerotiorum 555* grew well in media treated with no peptide. However, mycelial growth of this pathogen was progressively reduced with increasing concentrations of the peptide (Figure 1a). GMA4CG_V6 inhibited the growth of *S. sclerotiorum 555* in *vitro* with the half-maximal inhibitory concentration (EC_50_) value of ∼14 µM and the minimal inhibitory concentration (MIC) of 24 µM (Figure 1b). It also inhibited mycelial growth of two other *S. sclerotiorum* isolates, strains *1902* and *1922,* with similar potency (Figure 1c).

**Figure 1.**
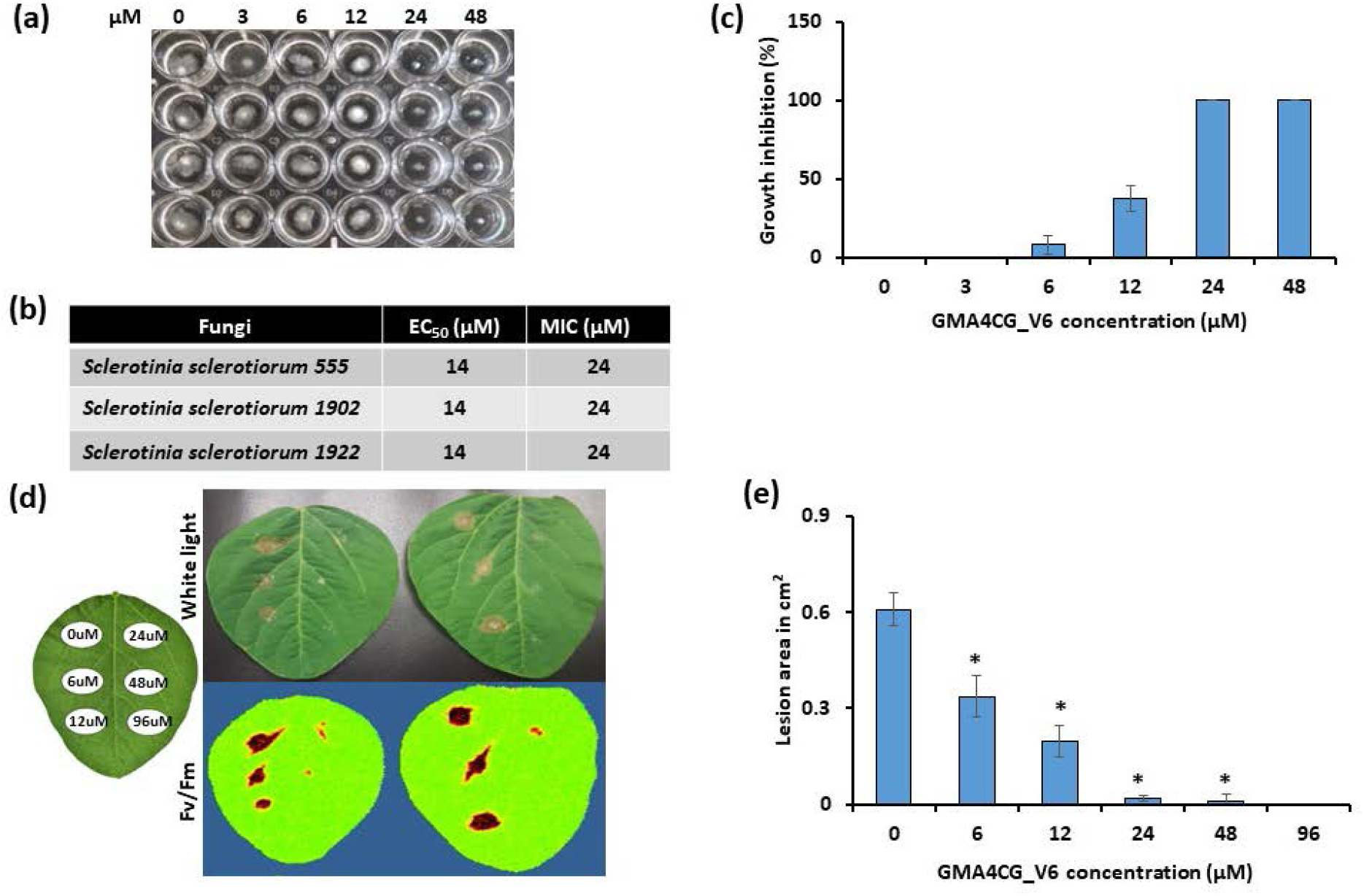
*In vitro* and semi-*in planta* antifungal activities of GMA4CG_V6 against *S. sclerotiorum 555* on soybean leaves. (**a**) Representative pictures showing the antifungal activity of GMA4CG_V6 against *S. sclerotiorum 555* in SFM media. (**b**) EC_50_ and MIC values of GMA4CG_V6 for *S. sclerotiorum 555.* **(c)** Inhibition of fungal growth at different concentrations of GMA4CG_V6. Each data point represents the average % growth from three replications and error bars represent the standard errors of the means between replicates. **(d)** Representative pictures (under white light and with CropReporter) showing the antifungal activity of GMA4CG_V6 against *S. sclerotiorum 555* on detached soybean leaves. **(e)** Relative disease lesion area following GMA4CG_V6 application on soybean leaf surface at different concentrations. Each data point represents the average lesion size from five leaves relative to control (no peptide, 0 µM), with error bars representing the standard errors of the means between replicates. Results were analyzed using ANOVA, followed by a Tukey’s post hoc test. * Indicates statistically significant compared to the control (*P* <0.05). This experiment was repeated many times with similar results.

### 3.2. Semi-in planta Antifungal Activity of GMA4CG_V6 against S. sclerotiorum 555

Next, we used a detached soybean leaf assay to determine if GMA4CG_V6 also provides antifungal activity *in planta.* Detached leaves of 3 to 4-week-old Williams 82 soybean plants were infected with 1 mm plugs of *S. sclerotiorum 555* followed by the application of a drop of GMA4CG_V6 at different concentrations on the plug. At 2 days post-infection (dpi), *S. sclerotiorum 555* caused large necrotic lesions on leaves treated with water alone. However, external application of GMA4CG_V6 on the leaf surface significantly decreased lesion sizes, with complete inhibition of disease observed at a concentration of 24 µM (Fig. 1d). The lesions caused by *S. sclerotiorum 555* infection were progressively reduced in size with increasing concentrations of GMA4CG_V6 as compared with lesions generated in absence of the peptide (Figure 1e). Following these results, we extended our investigation of the antifungal activity of GMA4CG_V6 against *S. sclerotiorum 555* on soybean pods and stems.

Detached soybean pods and stems from 6-week-old plants were infected with 1 mm mycelial plugs of *S. sclerotiorum 555* followed by the application on the plug of a drop of GMA4CG_V6 at different concentrations. At 3 dpi, peptide-treated pods and stems exhibited much slower lesion development rates compared with the water treated control pods and stems (Figure 2a-d). The progressively smaller disease lesions around the inoculation site were observed in both tissues with increasing concentrations of GMA4CG_V6 (Figure 2b, d). We observed that application of 48 and 96 µM GMA4CG_V6 onto stems and pods conferred full fungicidal activity with 100% protection against *S. sclerotiorum 555* at 3 dpi and also at 7dpi (Figure 2e, f). We conclude that external application of 24 to 96 µM GMA4CG_V6 to detached soybean leaves, pods, and stems significantly reduces disease lesions caused by infection of *S. sclerotiorum 555*.

**Figure 2.**
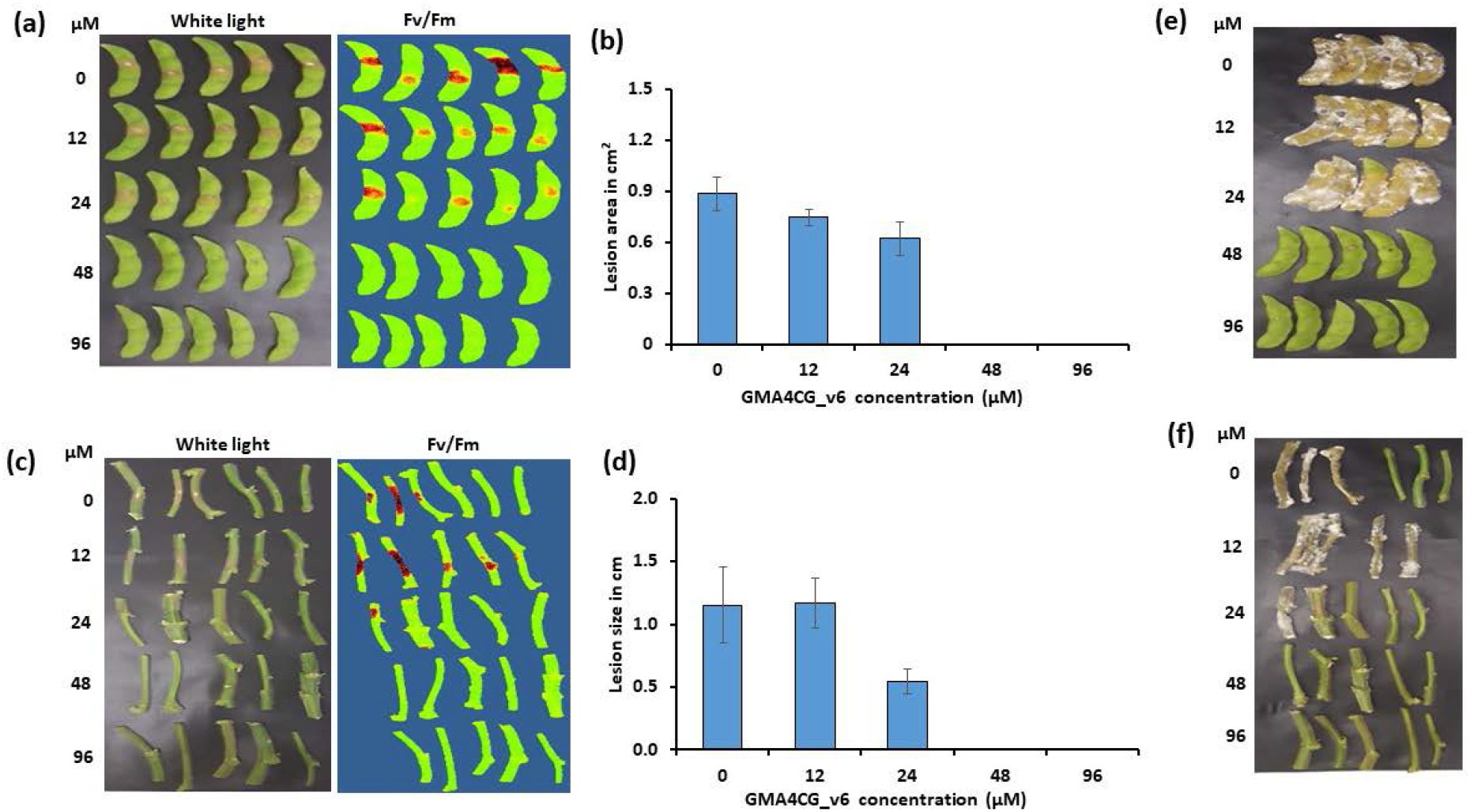
Semi-*in planta* antifungal activity of GMA4CG_V6 against *S. sclerotiorum 555* on soybean pods and stems. **(a)** Soybean pods detachment assay showing the effect of GMA4CG_V6 on disease development at 3 dpi. Representative pictures (under white light and with CropReporter). **(b)** Relative disease lesion area at 3dpi following the application of GMA4CG_V6 on the pods. **(c)** Soybean stems detachment assay showing the effect of GMA4CG_V6 on disease development at 3 dpi. **(d)** Relative lesion size following the application of GMA4CG_V6 on the stems surface at 3 dpi. Antifungal activities at 7dpi **(e, f).** For panel **(b)** and **(d)** each data point represents the average lesion area/size of five samples relative to control (no peptide, 0 µM), with error bars representing the standard errors of the means between replicates. This experiment was repeated three times with similar results.

### 3.3. Spray-application of GMA4CG_V6 Protects against White Mold Disease in Soybean and Nicotiana benthamiana

To test the potential of GMA4CG_V6 as a spray-on bio-fungicide for control of white mold disease on *N. benthamiana* and Willams 82 soybean plants, a whole plant assay was developed. Leaves of both species were spray-inoculated with a homogenized suspension of *S. sclerotiorum 555* mycelia followed by spray-application of GMA4CG_V6 at 24 and 48 µM onto the inoculated leaves. By 2 dpi, *N. benthamiana* leaves treated with GMA4CG_V6 at 24 to 48 µM concentrations showed much smaller lesions than water-treated plants (Figures 3a, b). Similarly, 24-48 µM GMA4CG_V6 reduced the size of necrotic lesions in soybean leaves at 2 dpi (Figure 3c, d). Thus, topical application of GMA4CG_V6 efficiently reduces white mold symptoms on both tobacco and soybean plants.

**Figure 3.**
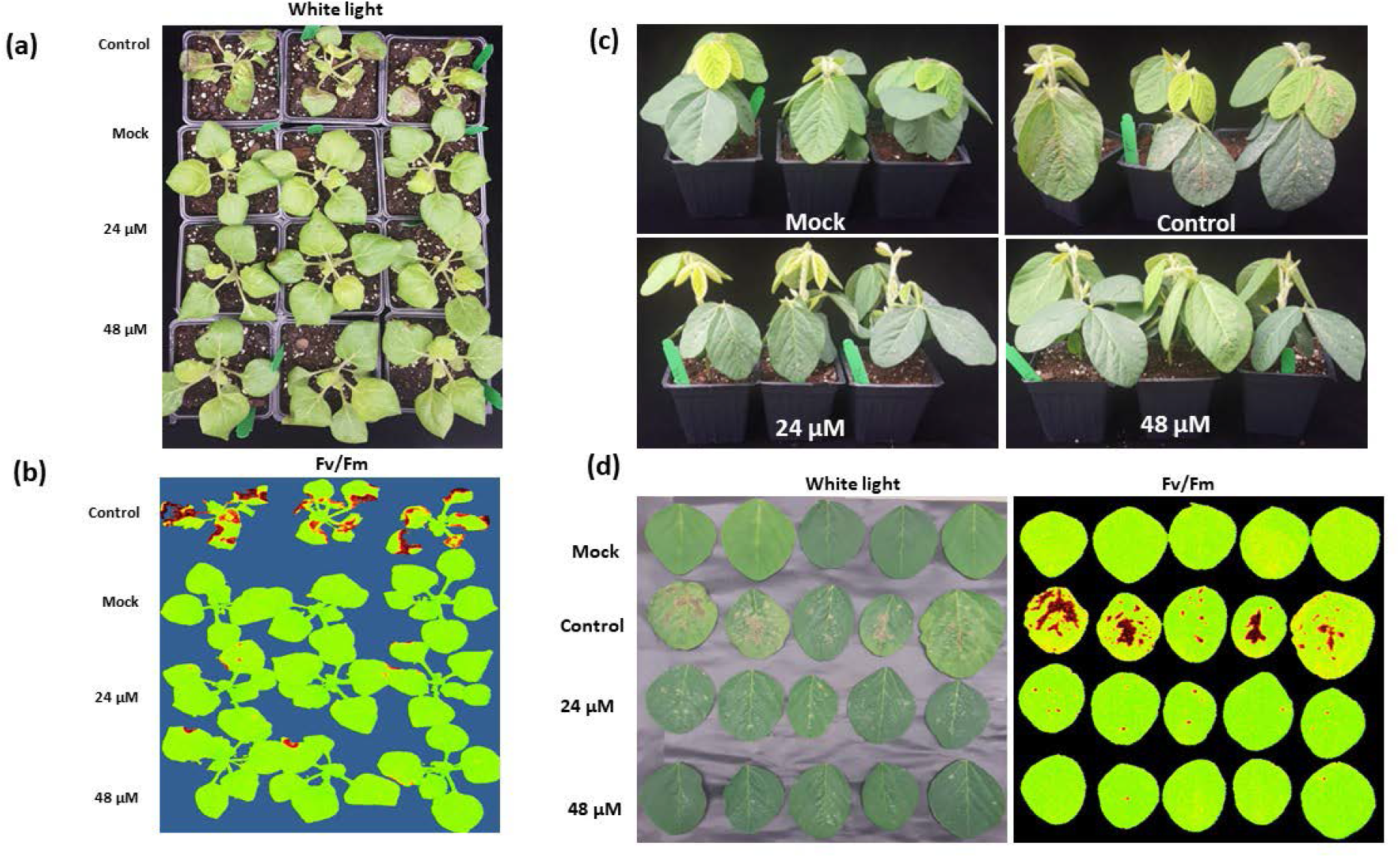
*In planta* antifungal activity of GMA4CG_V6 against *S. sclerotiorum 555*. **(a)** In-pot assay showing the antifungal activity of GMA4CG_V6 against *S. sclerotiorum 555* on tobacco leaves sprayed with *S. sclerotiorum 555* ground mycelia. **(b)** Representative pictures (CropReporter) showing the antifungal activity of GMA4CG_V6 against *S. sclerotiorum 555* on tobacco plants. **(c)** In-pot assay showing the antifungal activity of GMA4CG_V6 against *S. sclerotiorum 555* on soybean leaves sprayed with *S. sclerotiorum 555* ground mycelia. **(d)** Representative pictures (CropReporter) showing the antifungal activity of GMA4CG_V6 against *S. sclerotiorum* soybean leaves.

### 3.4. GMA4CG_V6 Inhibits Sclerotia Production in S. sclerotiorum 555

Since sclerotia play an important role in the disease cycle of *S. sclerotiorum*, we asked if GMA4CG_V6 affected sclerotia development. For these experiments, we used a sub-lethal concentration (12 µM) of GMA4CG_V6. For comparison, we also examined the effects of GMA4CG_V6_lin, a variant lacking a disulfide bond and GMA4CG_V6_Ala3, a variant with four-fold less antifungal activity against *B. cinerea* than GMA4CG_V6 [21]. The relative antifungal activity of these peptides against *S. sclerotiorum 555* was also determined to be similar to that against *B. cinerea* (Figure S1). Cinnamic acid treatment, known to inhibit sclerotia production, was used as a positive control. *S. sclerotiorum 555* mycelia growing on PDA plates were sprayed with water, cinnamic acid, or peptide on Day 1, Day 3 and Day 5 of growth. On Day 9, five sclerotia per plate had developed on the plates sprayed with water. However, only 1-2 sclerotia per plate had developed on plates sprayed with 12 µM GMA4CG_V6 and GMA4CG_V6_lin (Figure 4a, b, c). Interestingly, GMA4C_V6_Ala3, which has much reduced antifungal activity *in vitro,* failed to inhibit sclerotia production (Figure 4b, c), indicating a causal relationship between the *in vitro* antifungal activity of this peptide and its ability to inhibit sclerotia production. Spray application of GMA4CG_V6 peptide also significantly reduced sclerotia production on infected soybean pods (Figure 4d, e) Using confocal microscopy, we found that 12 µM GMA4CG_V6 prevented early development of sclerotia. In the absence of GMA4CG_V6, *S. sclerotiorum 555* hyphae initiated the production of new hyphal tips in clusters (sclerotia primordia) after two days of growth. In contrast, no early signs of sclerotia development were observed in the hyphae of *S. sclerotiorum 555* grown in the presence of 12 µM GMA4CG_V6 for two days (Figure 4f). Taken together, our results indicate that GMA4CG_V6 is able to inhibit the development of sclerotia in *S. sclerotiorum 555*.

**Figure 4.**
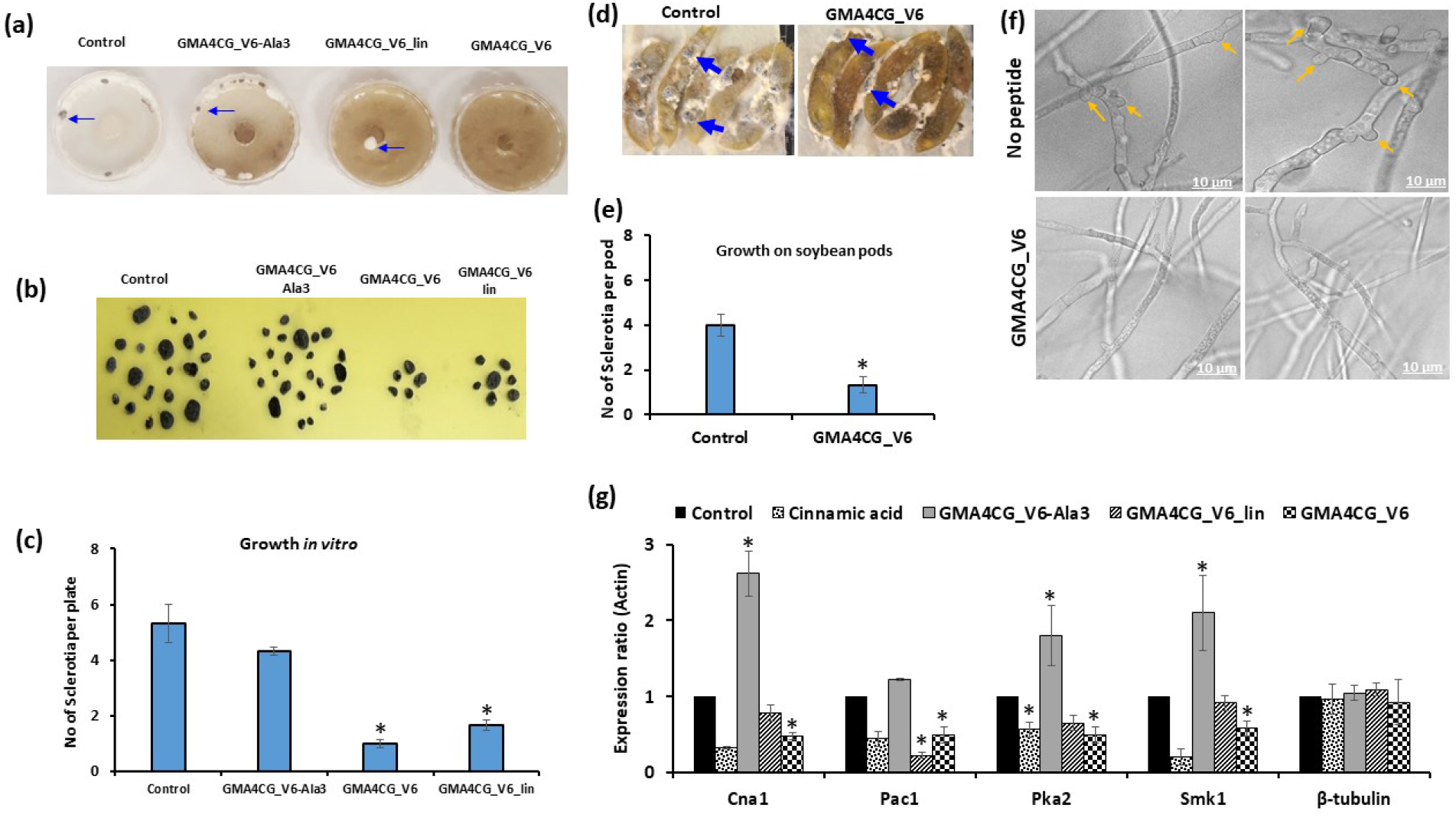
Effect of GMA4CG_V6 on sclerotia production by *S. sclerotiorum 555*. **(a)** Representative pictures showing the effect of various GMA4CG_V6 variants against sclerotia production *in vitro* after 14-day treatment. **(b)** Picture showing the total number of sclerotia produced 30 days following peptide treatment. Six plates were used for each treatment. (**c)** Average number of sclerotia per plate. Each data point represents the average of six plates, with error bars representing the standard errors of the means between replicates. Groups indicated with different letters are significantly different according to ANOVA with Tukey’s post hoc test. **(d)** Representative pictures showing the antifungal activity of GMA4CG_V6 against sclerotia production on soybean pods fourteen days post-treatment. (**e)** Average number of sclerotia per pod. Each data point represents the average of five pods, with error bars representing the standard errors of the means between replicates. **\*** indicates statistically significant difference (*P* <0.05) from Student’s t-test. **(f)** Confocal microscopy images showing the inhibition of sclerotia development by GMA4CG_V6 after two days of growth *in vitro*. Arrows showing the early development of sclerotia—growth of hyphal tips and dichotomous branching. **(g)** Effect of different GMA4CG_V6 variants on the expression level of genes involved in sclerotia production in *S. sclerotiorum 555.* Results were analyzed using a Student’s t-test. *indicates significant differences between control samples with *P* <0.05.

### 3.5. Effect of GMA4CG_V6 on the expression of S. sclerotiorum genes related to sclerotia production

We examined whether GMA4CG_V6 inhibits the expression of genes known to be essential for development of sclerotia [24] in *S. sclerotiorum 555*. Expression of *Cna1*, *Pac1*, *Pka2*, and *Smk1* genes was significantly reduced in mycelia sprayed with GMA4CG_V6 and GMA4CG_V6_lin (Figure 4g). Similar downregulation of the *Pac1* and *Smk1* gene expression was previously observed in mycelia treated with cinnamic acid [24]. We conclude that GMA4CG_V6 inhibits the development of sclerotia by downregulating the expression of key genes involved in their development. Surprisingly, expression of *Cna1*, *Pka2* and *Smk1* was significantly upregulated in mycelia sprayed with GMA4CG_V6_Ala3 peptide when compared with the expression of these genes in water-sprayed mycelia (Figure 4g).

### 3.6. GMA4CG_V6 permeabilizes the plasma membrane and is internalized into the cells of *S. sclerotiorum*

GMA4CG_V6 permeabilizes the plasma membrane and enters the cells of *B. cinerea* [21]. Since *B. cinerea* is far more sensitive to the antifungal activity of this peptide, we wondered if this peptide was also capable of permeabilizing the plasma membrane and traveling to the interior of the cells of *S. sclerotiorum 555*. The SYTOX Green (SG) uptake assay was used to test for plasma membrane integrity. SG uptake was visible in hyphal cells 30 min after challenge with a sub-lethal dose of 12 µM peptide (Figure 5a), indicating that membrane permeabilization contributes to the antifungal action of GMA4CG_V6. Next, we used TMR-labeled GMA4CG_V6 peptide to determine if the peptide was taken up by the cells of this pathogen. After 2-5 min treatment of *S. sclerotiorum 555* hyphae with 24 µM labeled peptide, it was internalized into fungal cells (Figure 5b). These observations suggest that membrane permeabilization and uptake into fungal cells are contributing factors to the MoA of GMA4CG_V6 in *S. sclerotiorum*.

**Figure 5.**
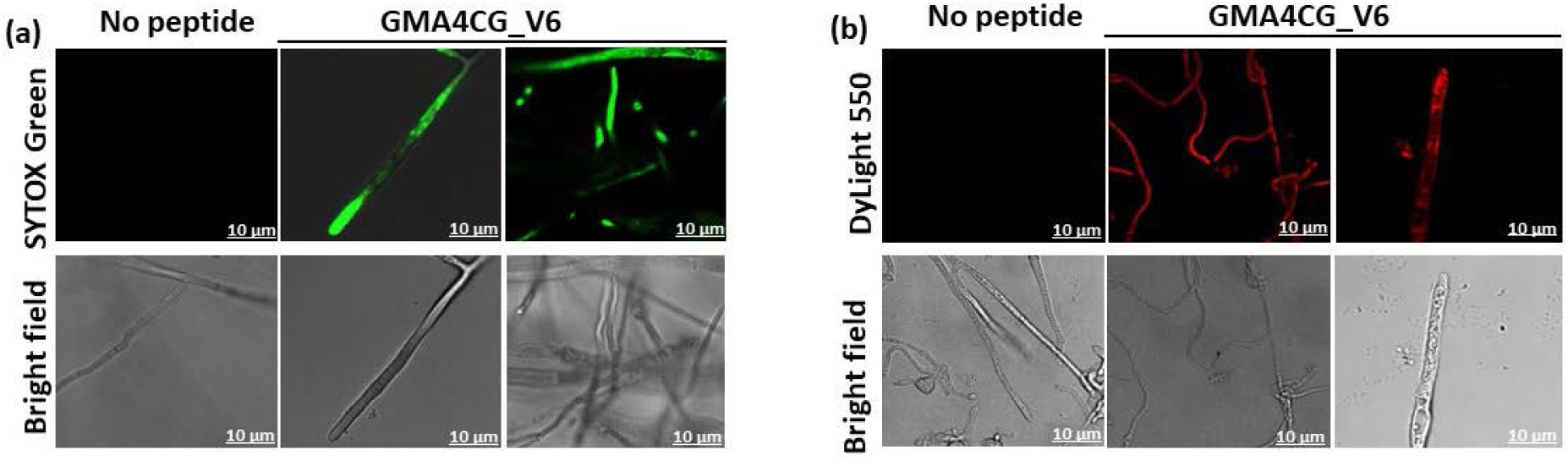
Membrane permeabilization activity and uptake of sub-lethal concentrations of GMA4CG_V6 by fungal cells. Confocal microscopy images and corresponding bright field images of SYTOX Green uptake in *S. sclerotiorum 555* hyphae treated with 12 μM GMA4CG_V6 for 30 min. (Scale bar, 10 μM). The intracellular localization of 24 µM TMR-labeled GMA4CG_V6 in *S. sclerotiorum 555*. The confocal microscope images were captured 2-5 min after GMA4CG_V6 challenge.

### 3.7. GMA4CG_V6 has no negative effect on the growth of soybean plants

Six weeks after GMA4CG_V6 treatment, we found no difference in the appearance of the leaves or the aerial growth and development of treated plants compared to control plants sprayed only with water (Figure 6). The root biomass and aerial biomass of the treated plants were similar to those of the control plants (Figure 6). Thus, GMA4CG_V6 has no discernible phytotoxicity.

**Figure 6.**
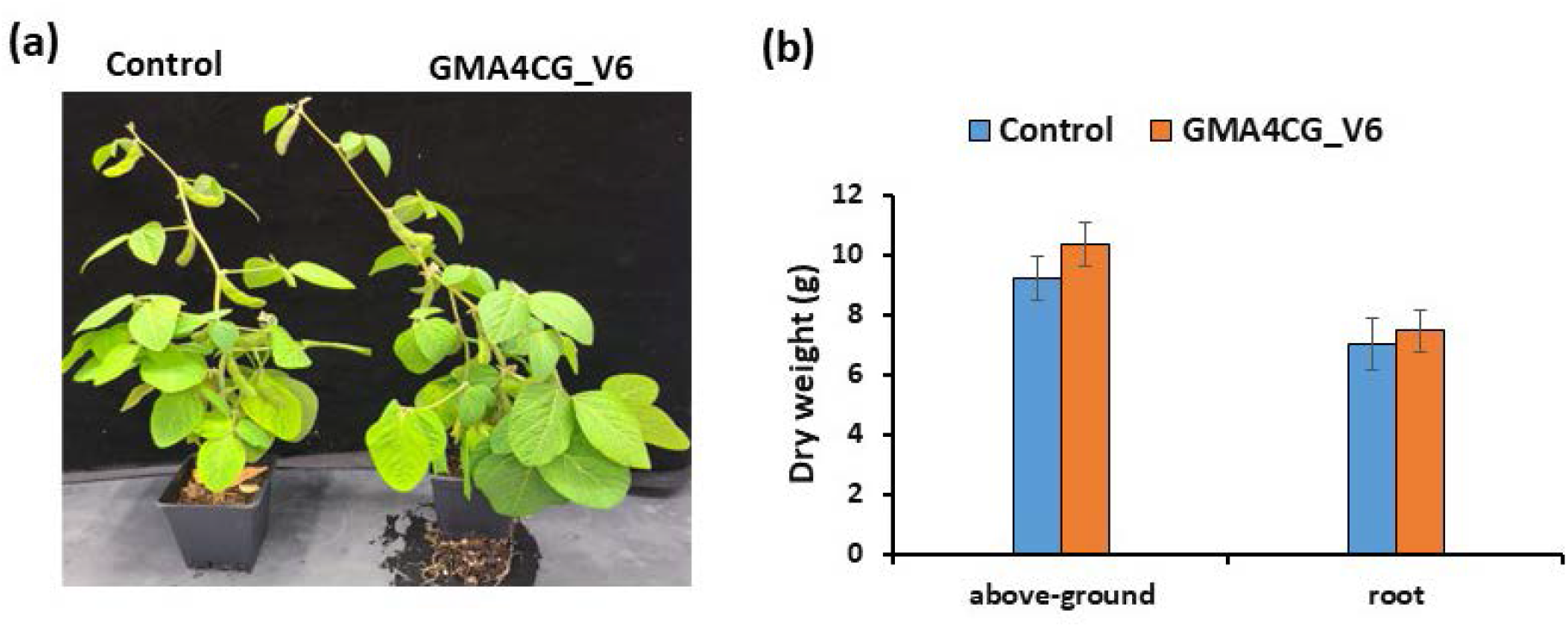
Effect of GMA4CG_V6 on soybean plant growth. **(a)** Representative picture showing no toxicity of GMA4CG_V6 treatment of soybean plants 45 days post treatment. **(b)** Above-ground and root biomass of plants treated with water or GMA4CG_V6 45-day post treatment. Each data point represents the average mass of five plants, with error bars representing the standard errors of the means between replicates.

## 4. Discussion

Small cysteine-rich antifungal peptides with multi-site MoA have strong potential for development as safe and sustainable bio-fungicides. In this study, we investigated a small 17-amino acid antifungal peptide, GMA4CG_V6, for its potential as a peptide-based bio-fungicide for management of white mold disease in soybean.

Until now, there has been a marked paucity of antifungal peptides active against *S. sclerotiorum*. Only one plant defensin, RsAFP1, has been reported to be active against this pathogen at 20 µg/ml (3.5 µM) [19]. However, potential of RsAFP1 to provide control of this pathogen *in planta* remains unclear as it was shown to reduce white mold symptoms in *B. napus* only in presence of a fungal chitinase [25].

GMA4CG_V6 is a variant of the GMA4CG peptide derived from the C-terminal sequence of an antifungal defensin, MtDef4. It is a pseudo-cyclic peptide with a disulfide bond connecting Cys4 and Cys17 and carries a net charge of +6 [21]. We recently reported that GMA4CG_V6 peptide exhibits both preventative and curative antifungal activity against *B. cinerea* when sprayed on tobacco and tomato plants [21]. In this study, we determined that GMA4CG_V6 reduced white mold symptoms when applied at the site of fungal inoculation on detached soybean leaves, pods, and stem after 2-3 dpi. No further white mold symptoms developed on pods and stems treated with 48 µM peptide even at 7 dpi. The lack of any symptoms on pods and stems suggests that this peptide has fungicidal activity against *S. sclerotiorum 555.* The pot-grown young soybean and tobacco plants sprayed with GMA4CG_V6 also showed marked reduction of the white mold symptoms when challenged with mycelial fragments of *S. sclerotiorum 555*, clearly demonstrating the potential of this peptide for development as a bio-fungicide.

Sclerotia are the main propagules for dispersal of *S. sclerotiorum* and play a major role in the disease cycle and infectivity of this pathogen in soybean [26]. The inhibition of sclerotial development by an antifungal agent could have a major positive impact on disrupting the disease cycle of this pathogen by reducing the sclerotial load in the soil. We observed that GMA4CG_V6 at sub-lethal concentrations was effective in inhibiting sclerotial development when sprayed on the mycelium of this fungus growing on petri plates as well as on detached soybean pods. The peptide was also able to reduce expression of a few genes known to be important for development of sclerotia [24]. Transcriptome analysis of the peptide-challenged mycelium will need to be performed to better understand the underlying mechanism of the peptide-induced inhibition of sclerotia development. To our knowledge, GMA4CG_V6 is the first example of an antifungal peptide known to inhibit sclerotia development. Similar inhibition of fungal growth, sclerotia development and gene expression was also reported for a natural product cinnamic acid against *S. sclerotiorum* [24].

GMA4CG_V6 exhibits stronger antifungal activity against *B. cinerea* with an MIC of only 1.5 µM [21]. In contrast, we found that this peptide was inhibitory against *S. sclerotiorum 555* only at a concentration of 24 µM. Antifungal potency of this peptide against two other *S. sclerotiorum* isolates was also similar. The difference in the response of *B. cinerea* and *S. sclerotiorum* is surprising given that these pathogens are closely related taxonomically belonging to the same family *Sclerotiniaceae* [23]. It is not clear what makes *S. sclerotiorum* much more tolerant to this antifungal peptide. In-depth MoA studies are needed to determine if *S. sclerotiorum* cell wall lacks high affinity binding sites for this peptide at a low concentration of 1.5 µM. It is also important to determine if significant differences in the mechanisms of internalization of this peptide and its intracellular targets exist between these two fungal pathogens.

Collectively, data presented in this study suggest that GMA4CG_V6 is effective at controlling *S. sclerotiorum 555* at multiple steps of the disease cycle, *i.e*., mycelial growth and sclerotial development, in its life cycle. Additionally, GMA4CG_V6 is likely a multi-site fungicide, as it not only disrupts the fungal plasma membrane but also binds to as-yet-unidentified intracellular targets in *S. sclerotiorum* (Figure 7).

**Figure 7.**
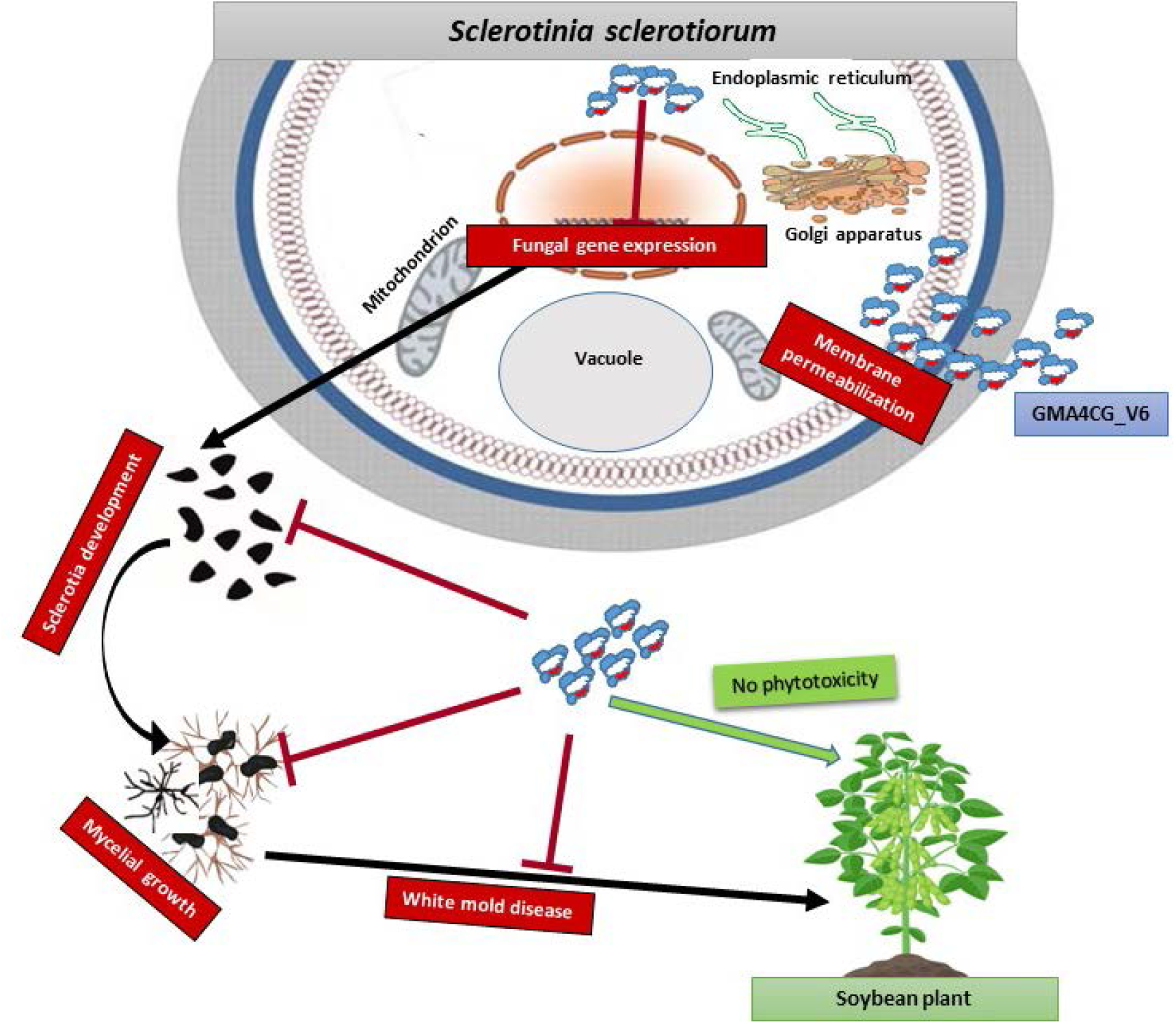
A proposed model illustrating the MoA of GMA4CG_V6 against *S. sclerotiorum 555*. GMA4CG_V6 sprayed on leaves infected with *S. sclerotiorum 555* first binds to the pathogen’s cell walls and the peptide starts to permeabilize the fungal plasma membrane. GMA4CG_V6 within the fungus can repress the expression of fungal genes involved in sclerotia development. The surface application of GMA4CG_V6 efficiently inhibits fungal growth and significantly reduces sclerotia development and white mold symptoms. Finally, soybean treatment with GMA4CG_V6 does not affect above- or below-ground biomass.

## 5. Conclusions

A short MtDef4 defensin-derived peptide GMA4CG_V6 inhibits the growth of *S. sclerotiorum in vitro*. Topical application of this peptide protects soybean plants from white mold and does not cause phytotoxicity. GMA4CG_V6 inhibits sclerotia production, permeabillizes the plasma membrane and gains entry into cells of this pathogen. This study demonstrates the potential of defensin-derived antifungal peptides as bio-fungicides.

## Acknowledgments

This work was supported by the National Sclerotinia Initiative grant 58-3060-1-026 awarded to D.S. We are very grateful to Dr. Sydney E. Everhart of the University of Nebraska for providing the *S. sclerotiorum* isolates used in this study. We acknowledge confocal microscopy support and guidance from the Advanced Bioimaging Laboratory (RRID:SCR_018951) at the Donald Danforth Plant Science Center. The Leica SP8-X confocal microscope used in this study was acquired through an NSF Major Research Instrumentation grant (DBI-1337680).

## Supporting information

**Table S1.**
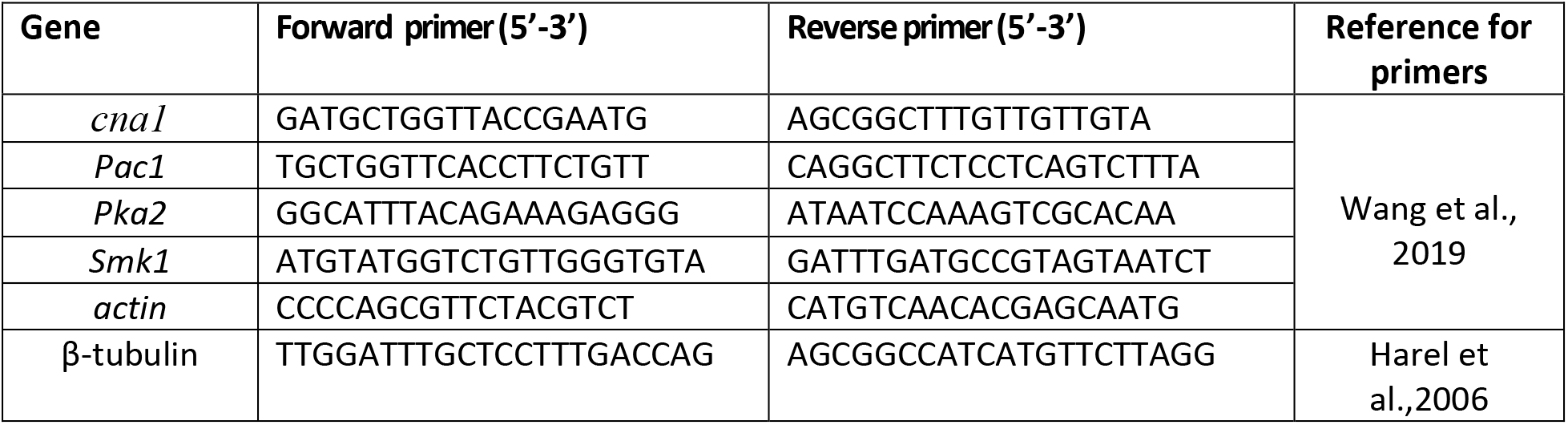
*S. sclerotiorum* gene-specific primers used for qPCR analysis of gene expression.

## Supporting Figure Legends

**Figure S1.**
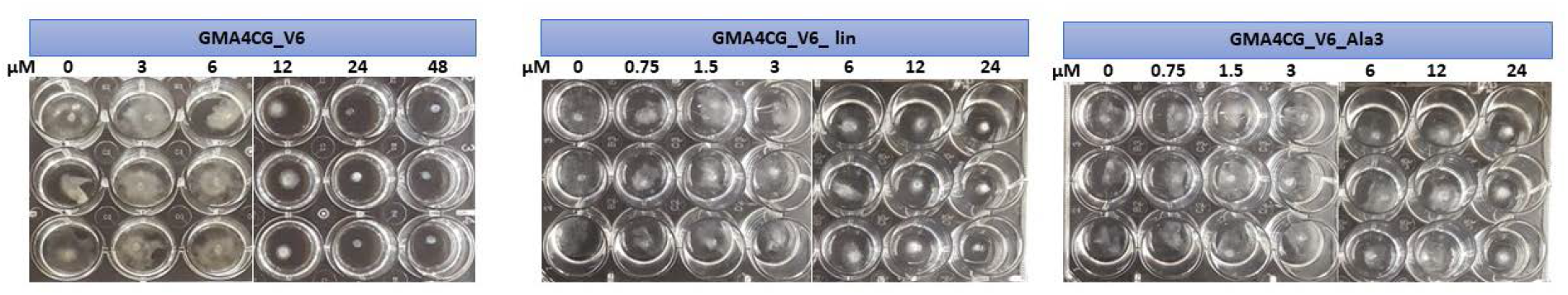
*In vitro* antifungal activity of GMA4CG_V6 variants against *S. sclerotiorum 555*. Representative pictures showing the antifungal activity of GMA4CG_V6, GMA4CG_V6_lin and GMA4CG_V6_Al3 against *S. sclerotiorum 555, 1902* and *1922* in SFM media.

